# Chalkophomycin Biosynthesis Revealing Unique Enzyme Architecture for a Hybrid Nonribosomal Peptide Synthetase and Polyketide Synthase

**DOI:** 10.1101/2024.03.14.584926

**Authors:** Long Yang, Liwei Yi, Bang Gong, Lili Chen, Miao Li, Xiangcheng Zhu, Yanwen Duan, Yong Huang

**Affiliations:** Department of Immunology, School of Basic Medical Sciences, Anhui Medical University, Hefei 230032, China; Institute of Health and Medicine, Hefei Comprehensive National Science Center, Hefei, Anhui, 230093, China; Xiangya International Academy of Translational Medicine, Central South University, Changsha, Hunan, 410013, China; The Affiliated Nanhua Hospital, Department of Pharmacy, Hengyang Medical School, University of South China, Hengyang, Hunan, 421001, China; College of Pharmacy, Hunan Vocational College of Science and Technology, Changsha, 410004, China; Hunan Engineering Research Center of Combinatorial Biosynthesis and Natural Product Drug Discover, Changsha, Hunan, 410011, China; National Engineering Research Center of Combinatorial Biosynthesis for Drug Discovery, Changsha, Hunan, 410011, China

**Author notes:** Equal contribution (Long Yang, Liwei Yi).

**Keywords:** chalkophore, copper, peptide, hybrid NRPS/PKS, chain release mechanism, reductase domain (R^0^)

## Abstract

Chalkophomycin is a novel chalkophore with antibiotic activities isolated from *Streptomyces* sp. CB00271, while its potential in studying cellular copper homeostasis makes it an important probe and drug lead. The constellation of *N*-hydroxylpyrrole, 2*H*-oxazoline, diazeniumdiolate, and methoxypyrrolinone functional groups into one compact molecular architecture capable to coordinate cupric ion draws interest to unprecedented enzymology responsible for chalkophomycin biosynthesis. To elucidate the biosynthetic machinery for chalkophomycin production, the *chm* biosynthetic gene cluster from S. sp. CB00271 was identified, and its involvement in chalkophomycin biosynthesis was confirmed by gene replacement. The *chm* cluster was localized to a ∼31 kb DNA region, consisting of 19 open reading frames that encode five non-ribosomal peptide synthetase (ChmHIJLO), one modular polyketide synthases (ChmP), six tailoring enzymes (ChmFGMNQR), two regulatory proteins (ChmAB), and four resistance proteins (ChmA′CDE). A model for chalkophomycin biosynthesis is proposed based on functional assignments from sequence analysis and structure modelling, and is further supported by analogy to over 100 *chm*-type gene clusters in public databases. Our studies thus set the stage to fully investigate chalkophomycin biosynthesis and to engineer chalkophomycin analogues through a synthetic biology approach.

## 1. Introduction

Chalkophomycin, originally isolated from *Streptomyces* sp. CB00271 in 2021, is an unprecedented copper(II)-binding metallophore (Figure 1A) [1]. The gross structure of chalkophomycin was deduced by single-crystal X-ray analysis, and its X-ray photoelectron spectroscopy analysis revealed that Cu(II) is the majority of copper species. The structure of the copper-less apo-chalkophomycin was established by spectroscopic analysis, and its absolute stereochemistry was based on the similar CD spectra with chalkophomycin. These analysis revealed that Cu(II) in chalkophomycin is coordinated to *N*-hydroxylpyrrole, 2*H*-oxazoline, and diazeniumdiolate from a methoxypyrrolinone ring. These respective functional groups in chalkophomycin have been found in dozens of natural products, e.g, glycerinopyrin, hormaomycins, and surugapyrroles (for *N*-hydroxylpyrrole), coelibactin, mycobactin, and aerucyclamides (for 2*H*-oxazoline), alanosine, fragin, and gramibactin (for diazeniumdiolate), althiomycin, dolastatin 15, and malyngamide A (for methoxypyrrolinone) [2–16]. However, compact integration of these distinct elements into one molecular architecture capable to coordinate cupric ion highlights nature’s unique design strategy for chalkophores, emerging family of natural products responsible for microbial copper homeostasis [17].

**Figure 1.**
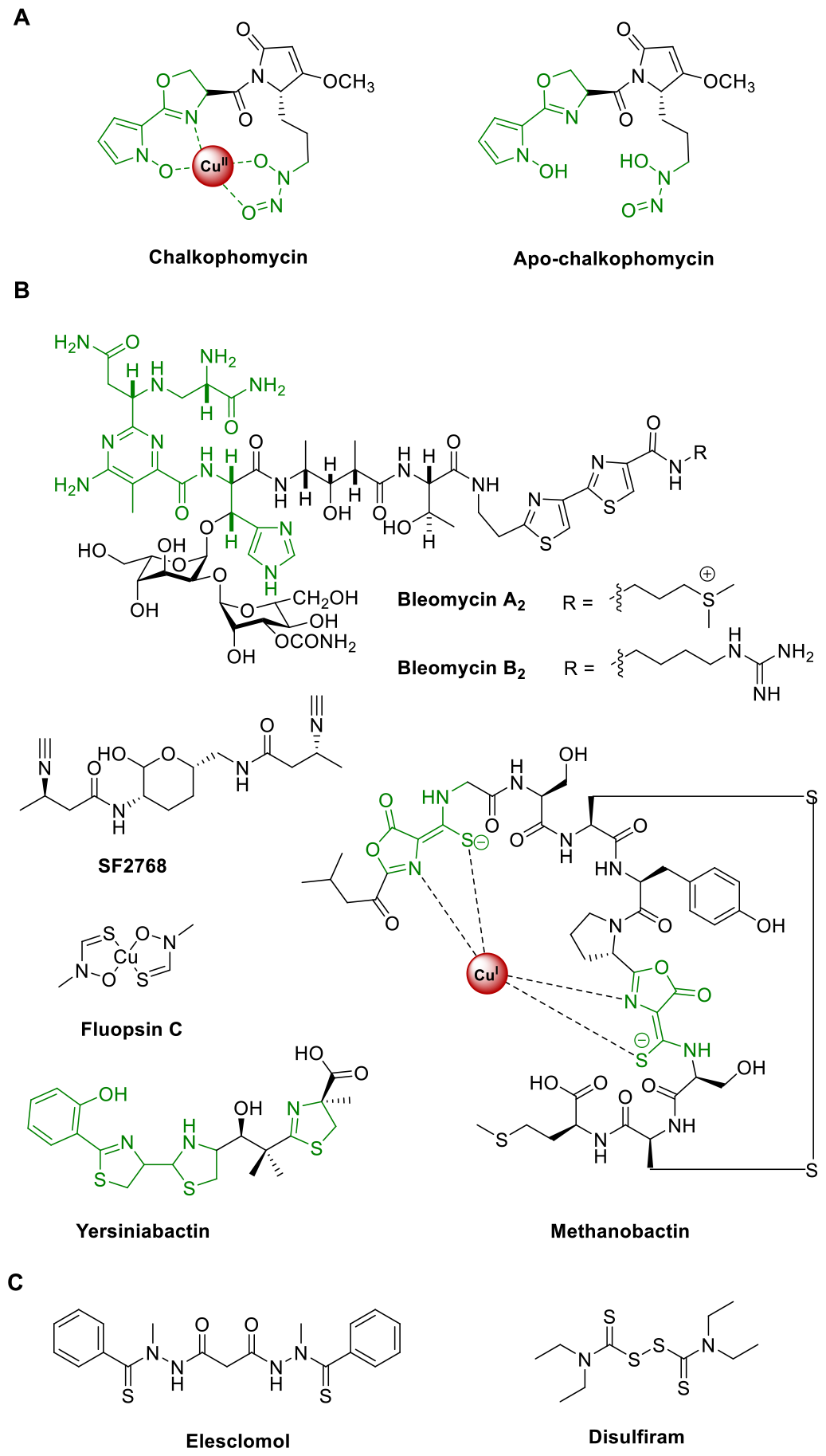
Representative natural and synthetic chalkophores.

In an analogy to siderophores for iron homeostasis, there is growing interest to study the biological function and biosynthesis of chalkophores [18–26]. Chief among them are members of the methanobactin family, which were first identified from methane-oxidizing bacteria. These methanotrophic bacteria produce abundant amount of copper enzyme named particulate methane monooxygenase, which catalyzes the aerobic oxidation of methane and plays indispensable role in global carbon cycle. Recent genome mining efforts have revealed that some other bacteria may also produce methanobactins for copper acquisition [19]. A wider range of bacteria can produce chalkophores, including bleomycin, yersiniabactin, SF2768, and xanthocillins (Figure 1B). Interestingly, yersiniabactin was initially discovered as a siderophore, but its noncanonical role for copper and other non-iron metal ions uptake was recently discovered in pathogenic *Enterobacteriaceae*, supporting intricate interactions between host and pathogens mediated by natural products and transition metal ions [27–28]. Copper can not only serve as an active site cofactor for certain proteins, e.g. “blue” copper proteins, and the aforementioned particulate methane monooxygenase, but also regulate protein function allosterically in signaling pathways in cancer, fatty liver disease, neurodegeneration, and obesity [29]. Therefore, methanobactins have been used in the treatment of acute Wilson’s disease in a WD rat model to alleviate copper overload, since excess copper causes hepatocyte death [30]. In addition, cuproptosis, a new form of cell death targeting lipoylated TCA cycle proteins, was recently discovered using elesclomol, a synthesized chalkophore (Figure 1C) [31]. Combination treatment with copper and disulfiram, an old drug against alcohol-abuse, also showed promise to induce cancer cell cuproptosis [32]. Taken together, these copper-binding molecules represent interesting drug leads and powerful small molecule probes to elucidate the roles of copper-signaling pathways.

The purpose of our study was to discover and characterize the chalkophomycin biosynthetic gene cluster (*chm*). The long-term goal of our program focused on chalkophomycin is to develop novel probes for cuproptosis and cuproplasia, and potential drug leads for the treatment of cancer and Wilson’s diseases. We report here on (i) the discovery and genetic characterization of the *chm* gene cluster in *S*. sp. CB00271; (ii) bioinformatics analysis of the *chm* cluster and a biosynthetic proposal for chalkophomycin biosynthesis involving a hybrid nonribosomal peptide synthetase and polyketide synthase (NRPS/PKS); (iii) genome mining of the *chm* pathway revealing its global distribution in a wide range of actinomycetes. Our study now enables rapid access to chalkophomycin gene clusters, as well as genome mining of individual biosynthetic enzymes for the formation of *N*-hydroxylpyrrole, 2*H*-oxazoline, diazeniumdiolate, and methoxypyrrolinone. The stage is now set for a synthetic biology approach to engineer chalkophomycin analogues as small molecule probes, drug leads, and potential chiral transition metal catalysts.

## 2. Materials and Methods

### 2.1. General experimental procedures

All chemical and biological reagents were purchased from commercial sources unless otherwise specified. Chalkophomycin and crude extracts were analyzed by Waters E2695 HPLC system equipped with a Welch AQ–C18 column (5 μm, 250 × 4.6 mm, Welch Materials Inc.) and detected with a photodiode array detector. Genomic DNA was isolated following standard protocols [33]. Plasmid DNA was extracted and purified using a PM0201 kit (Tsingke Biotech. Co.). The restriction endonucleases were purchased from New England Biolabs. DNA manipulation was based on standard procedures, including restriction endonuclease digestion and transformation.

### 2.2. Strains, plasmids, and culture conditions

*Streptomyces* sp. CB00271 was preserved in our lab. For sporulation, all strains were grown at 30°C on R2A solid medium. *Escherichia coli* DH5α and S17–1 were used for clon– ing and intergeneric conjugation respectively, and all were cultured with Luria–Bertani medium. All conjugants were grown on mannitol soya flour solid medium containing 10 mM MgCl_2_. For the cultivation of corresponding mutants, antibiotics including 50 mg/L apramycin, 25 mg/L thiostrepton, and 40 mg/L nalidixic acid were supplemented accordingly. All applied media were described in supporting information. All strains and plasmids were listed in Table S1.

### 2.3. *Fermentation production and HPLC analysis* of *chalkophomycin*

The spores of *Streptomyces* sp. CB00271 and its mutant strains were inoculated into Erlenmeyer flasks (250 mL) containing 50 mL of tryptic soy broth medium (1.7% tryptone, 0.3% soya peptone, 0.25% dextrose, 0.5% NaCl, 0.25% K_2_HPO4, pH 7.3) at 28 °C on a shaker at 230 rpm for 24∼48 h, with or without the addition of antibiotics. Then ∼10% (v/v) seed cultures were transferred into 50 mL production medium (2% soluble starch, 2% soy bean flour, 0.05% KH_2_PO_4_, 0.025% MgSO_4_) in 250 mL Erlenmeyer flasks. The pH of the production medium was adjusted to 7.0, followed by the addition of 0.5% (w/v) CaCO_3_ and 8.0% (v/v) macroporous resins DA201-H (Jiangsu Su Qing Water Treatment Engineering Group Co., Ltd., Jiangyin, China). These *Streptomyces* strains were then cultured for 7 days on a shaker at 230 rpm/28 °C.

For HPLC analysis of chalkophomycin production, the mobile phase included buffer A (ultrapure H_2_O containing 0.1% HCO_2_H) and buffer B (chromatographic grade CH_3_CN containing 0.1% HCO_2_H). A linear gradient program (95% buffer A for 2 min; 95% buffer A to 5% buffer A for 20 min; 5% buffer A for 2 min; 5% buffer A to 95% buffer A for 1 min, followed by 95% buffer A for 2 min) was applied at a flow rate of 1 mL/min.

### 2.4. Gene replacement of chmO in S. sp. CB00271

A pOJ260-based plasmid pXY5001 was constructed to generate the *ΔchmO* gene replacement mutant in *S*. sp. CB00271 via a double-crossover homologous recombination. To inactivate *chmO*, a 546-bp fragment of the *chmO* gene was replaced with the thiostrepton resistance gene with a *kasOp*^*\**^ promoter using the In-Fusion cloning kit (Tsingke, China), and the mutated *chmO* gene was cloned into pOJ260 between the *Hind*III and *Xba*I restriction sites. This plasmid was introduced into *Streptomyces* sp. CB00271 by conjugation and selected for thiostrepton resistance and apramycin-sensitive phenotype to isolate the desired double-crossover mutant strains. The PCR primers are shown in Table S2.

### 2.5. Structural analysis of the ChmP_R^0^ domains in S. sp. CB00271

The ChmP_R^0^ domain in *S*. sp. CB00271 was predicted using AlphaFold2 [34]. Molecular docking was performed by AutoDock Vina the predicted model of ChmP_R^0^ domains [35]. AutoDock Tools (The Scripps Research Institute, La Jolla, California, USA) was used to prepare the ligands and receptor as pdbqt files after removing water, adding polar hydrogen atoms and Gasteiger charges, respectively. The docking grid box size used was adjusted accordingly to encompass the NADP interaction site. Other default parameters were used. The best docking pose (most stable) was selected for binding mode comparison. The ligand-protein interaction structures were generated in PyMol (The PyMOL Molecular Graphics System, Version 3.0 Schrödinger, Inc.) [36].

### 2.6. Gene cluster similarity network analysis of chm genes in public databases

In order to identify homologous *chm* gene clusters from GenBank, cblaster (version 1.3) was used to search for similar gene clusters from the nonredundant database in GenBank with 50% sequence identity cutoff and the default parameters of 20,000 Max intergenic gap [37]. These identified *chm*-like genes contain at least eight homologous genes from the identified *chm* gene cluster in *S*. sp. CB00271. In addition, a Blastn search with *chmP* as the probe (50% identity cutoff) was performed [38]. The identified gene clusters were manually checked to remove duplicated gene clusters. Similarly, similar *chm* gene clusters were also identified by BlastP search using ChmP from the Natural Product Discovery Center actinomycete genome database from University of Florida, Scripps Research, using 50% sequence identity cutoff [39]. BIG-SCAPE (version 1.1.5) was used to analyze these gene clusters with a default parameter cutoff of 0.3 [40]. The resulting data were visualized using the organic layout in Cytoscape (version 3.10) [41]. Clinker 0.0.28 is used to generate cluster comparisons when running in the Basic pipeline [42].

## 3. Results

### 3.1. Discovery and genetic characterization of the chm gene cluster in S. sp. CB00271

The *chm* gene cluster was discovered by genome mining of putative biosynthetic genes in *S*. sp. CB00271 responsible for diazeniumdiolate and methoxypyrrolinone formation (Figure 2). There are over 300 nitrogen-nitrogen bond-containing natural products discovered, which bear a variety of important functional groups, e.g. diazo, hydrozones, pyrazole, and diazeniumdiolate [43]. Pioneering studies of diazeniumdiolate biosynthesis in streptozotocin, l-alanosine, and fragin/valdiazen revealed multiple unique enzymes *en route* for *N*-*N* construction and further morphing from amino acid precursors, including SnzE/SznF, AlnDEFGLMN, HamACED/HamACEDG [3, 7–9, 44–45]. In particular, Hertweck and co-workers recently discovered GrbED and their homologs responsible for biosynthesis of l-graminine, an unnatural amino acid found in a handful of cyclic or linear peptides, such as gramibactin, gladiobactin, JBIR-141/JBIR-142,megapolibactins, plantaribactin, and trinickiabactins [10–11]. These l-graminine-containing peptides are ubiquitously assembled by NRPSs, during which l-graminine is activated by specific adenylation domains and loaded onto the thiolation domain in NRPS assembly line. We therefore hypothesized that the diazeniumdiolate biosynthesis and loading in chalkophomycin is likely to follow the similar logic for the biosynthesis of gramibactin and alike, in which a similar analogous enzyme pair of GrbED should be present in *chm* gene cluster. Therefore, using GrbD and GrbE as query sequences, we identified three sets of GrbD and GrbE homologous genes from the genome of *S*. sp. CB00271 (Figure S1). Sequence alignment revealed that they share 33%–35% sequence identity with GrbE, and 38% sequence identity with GrbD, respectively. AntiSMASH-based analysis predicted that they are located in the flanking regions of several NRPSs in the genome of *S*. sp. CB00271, similar to the gramibactin gene cluster in *Paraburkholderia graminis* [10, 46].

**Figure 2.**
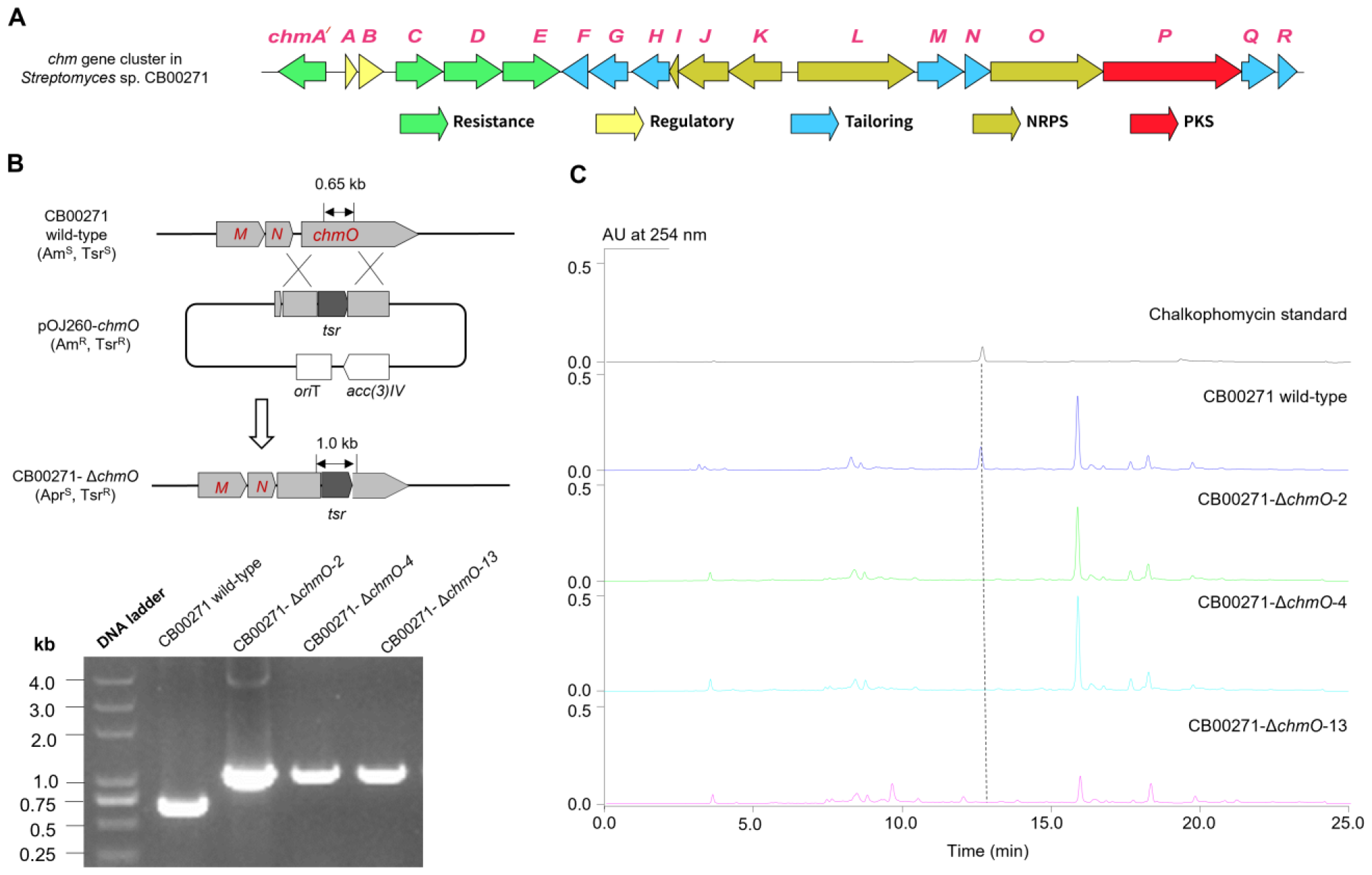
Identification and confirmation of the *chm* gene cluster. (**A**) The *chm* gene cluster contains a total of 19 ORFs from *chmA′* to *chmR*; (**B**) Gene replacement of *chmO* NRPS gene by a thiostrepton-resistant gene; (**C**) HPLC analysis revealed that the three *S*. sp. CB00271::Δ*chmO* mutants all abolished the production of chalkophomycin, in comparison to the wild-type strain.

In addition, the methoxypyrrolinone moiety in chalkophomycin is also found in several other natural products, including althiomycin,dolastatin 15,sintokamide A,malyngamide A, and mirabimide E [13–14, 47–48]. A malonyl-specific PKS module and a standalone *O*-methyltransferase were proposed for methoxypyrrolinone biosynthesis in althiomycin (Figure 2b) [13–14]. Therefore, close examination of the flanking regions of GrbE and GrbE homologs revealed that one cluster has a predicted PKS (WP_073800092.1) and an O-methyltransferase (WP_073800093.1). This PKS shows 42% sequence identity with module six of AlmB for althiomycin, while the *O*-methyltransferase shows 42% sequence identity with PokM3, responsible for *O*-methylation in pikromycin biosynthesis (Figure S2) [49]. Therefore, this gene cluster was named the chalkophomycin biosynthetic gene cluster (*chm*). In contrast, there are no such PKS and *O*-methyltransferase genes in the other two gene clusters.

The overall GC content of *chm* gene cluster is 72%, similar to other *Streptomyces* DNA. Bioinformatic analysis of *chm* cluster revealed 19 open reading frames (ORFs) (Figure 2b). Comparison of the deduced gene products from the *chm* gene cluster with proteins of known functions in the database facilitated the functional assignment of individual ORFs (Table 1). While this manuscript was in preparation, the same *chm* cluster from *Streptomyces* sp. CB00271 was reported to be responsible for chalkophomycin biosynthesis based on genome mining of SznF for streptozotocin biosynthesis, albeit without *chmA′* and *chmR*; gene replacement of this cluster however was not performed [50].

**Table 1.** Deduced Functions of Open Reading Frames in the Chalkophomycin Biosynthetic Gene Cluster.

| gene | size | putative function | Protein homologue | identity%<br>/similarity % |
| --- | --- | --- | --- | --- |
| *Orf(-2)* | 293 | diacylglycerol kinase | WP_011029127.1 | 60/67 |
| *Orf(-1)* | 427 | Adenylosuccinate synthase | 4M0G_A | 53/67 |
| *ChmA'* | 484 | MFS transporter | EfpA (ALB20045) | 34/53 |
| *ChmA* | 119 | regulatory LuxR family protein | RimR2 (QAS68949) | 35/59 |
| *ChmB* | 248 | TetR/AcrR-like transcription regulators | SCO1718 (CAB50933) | 32/40 |
| *ChmC* | 478 | MFS transporter | EfpA (ALB20045) | 32/51 |
| *ChmD* | 595 | ABC transporter | SCO7689 (CAC17506) | 48/62 |
| *ChmE* | 582 | ABC transporter | BDD77077 | 46/60 |
| *ChmF* | 264 | proline iminopeptidase | AlmF (CCA29204) | 24/37 |
| *ChmG* | 395 | acyl-CoA/acyl-ACP dehydrogenase | TdaE (WP_014881725.1) | 25/42 |
| *ChmH* | 383 | L-prolyl-PCP dehydrogenase | CloN3 AAN65232) | 31/41 |
| *ChmI* | 92 | peptidyl carrier protein | CloN5 AAN65234) | 26/54 |
| *ChmJ* | 517 | adenylation protein | CloN4 ( AAN65233) | 31/48 |
| *ChmK* | 543 | adenylation protein | EntE (CAD6013920) | 40/56 |
| *ChmL* | 1189 | non-ribosomal peptide synthetase | AlmA (CCA29202) | 33/48 |
| *ChmM* | 469 | L-graminine biosynthesis | GrbE (WP_006051176.1) | 35/48 |
| *ChmN* | 264 | L-graminine biosynthesis | GrbD (WP_006051175.1) | 37/51 |
| *ChmO* | 1152 | non-ribosomal peptide synthetase | AlmA (CCA29202) | 37/53 |
| *ChmP* | 1414 | type I polyketide synthase | AlmB (CCA29203) | 42/54 |
| *ChmQ* | 339 | O-methyltransferase | PokMT3 (ACN648470) | 36/50 |
| *ChmR* | 191 | flavin reductase | VlmR (AAC45645) | 32/50 |
| *Orf(+1)* | 273 | chitinase | Chitinase C (1WVU_A) | 58/75 |
| *Orf(+2)* | 294 | chitinase | Chitinase C (1WVU_A) | 97/98 |
| *Orf(+3)* | 371 | DNA polymerase III subunit beta | 5AH2_A | 56/58 |

In order to study if *chmO* encodes an NRPS from the *chm* gene cluster is involved in the biosynthesis of chalkophomycin in *S*. sp. CB00271, a 546-bp DNA fragment inside of *chmO* was replaced by a mutant copy in which *chmO* was disrupted by the thiostrepton-resistant gene with a *kas*Op^*^ promoter (Figure 2b) [51]. The gene replacement of *chmO* completely abolished the production of chalkophomycin in *S*. sp. CB00271 (Figure 2c). This result suggests that the identified *chm* gene cluster in *S*. sp. CB00271 is responsible for chalkophomycin biosynthesis.

### 3.2. Bioinformatics analysis of the chm cluster in S. sp. CB00271 revealed a hybrid NRPS/PKS for chalkophomycin biosynthesis

#### 3.2.1. Overview of the *chm* gene cluster

The *chm* gene cluster encompasses 19 ORFs designated *chmA’* to *chmR* (Figure 2a and Table 1). These biosynthetic genes encode NRPSs (ChmI, ChmJ, ChmK, ChmL, ChmO), PKS (ChmP), and other tailoring enzymes (ChmF, ChmG, ChmH, ChmM, ChmN, ChmQ, ChmR); Among them, two are regulatory genes (ChmA and ChmB), and four are resistance genes (ChmA*’*, ChmC, ChmD, and ChmE).

#### 3.2.2. Biosynthesis of NRPS precursor *N*-hydroxylpyrrole and l-graminine

Although pyrrole is found in a number of natural products biosynthesized by NRPSs, including clorobiocin, cloumermycin A1, plyoluteorin, prodigiosin, and undecylprodigiosin, *N*-hydroxylpyrrole building block is rare in nature. Considering that the formation of some pyrroles is catalyzed by a four-electron, two-step process from proteinogenic amino acid l-proline mediated by FAD-dependent reductases [52–54], biosynthesis of *N*-hydroxylpyrrole may adapt a similar route to generate a *N*-pyrrolyl-2-thioester-peptidyl carrier protein (PCP), followed by *N*-oxidation *en route* to *N*-hydroxylpyrrolyl-2-carboxyl-S-PCP. Careful examination of *chm* gene cluster revealed a small “*N*-hydroxylpyrrole” gene cassette containing *chmGHIJK* putatively responsible for *N*-hydroxylpyrrole biosynthesis in chalkophomycin (Figure 3). The identified genes were: (1) a free-standing 50-kDa l-proline specific adenylation (A) domain ChmJ responsible for l-proline activation to from l-prolyl-AMP, (2) The free-standing PCP ChmI would be loaded with l-prolyl-AMP to form l*-*pyrolyl-*S*-ChmI, (3) a predicated flavoprotein ChmH is presumably responsible for oxidizing pyrolyl-*S*-ChmI to pyrrolyl-*S*-ChmI, since it resembles CloN3, an l-prolyl-PCP dehydrogenase for pyrrole biosynthesis in antibiotic clorobiocin [53]; (4) another flavoprotein ChmG may catalyze pyrrolyl-*S*-ChmI oxidation to form *N*-pyrrolyl-*S*-ChmI. ChmF is predicated to be a proline iminopeptidase, sharing 24% and 37% sequence identity and similarity to AlmF in althiomycin biosynthesis in *M. xanthus* DK897, respectively, which is proposed for its methoxylpyrrolinone formation. However, the role of ChmF in *N*-hydroxylpyrrole formation or methoxylpyrrolinone biosynthesis in chalkophomycin remains to be determined.

**Figure 3.**
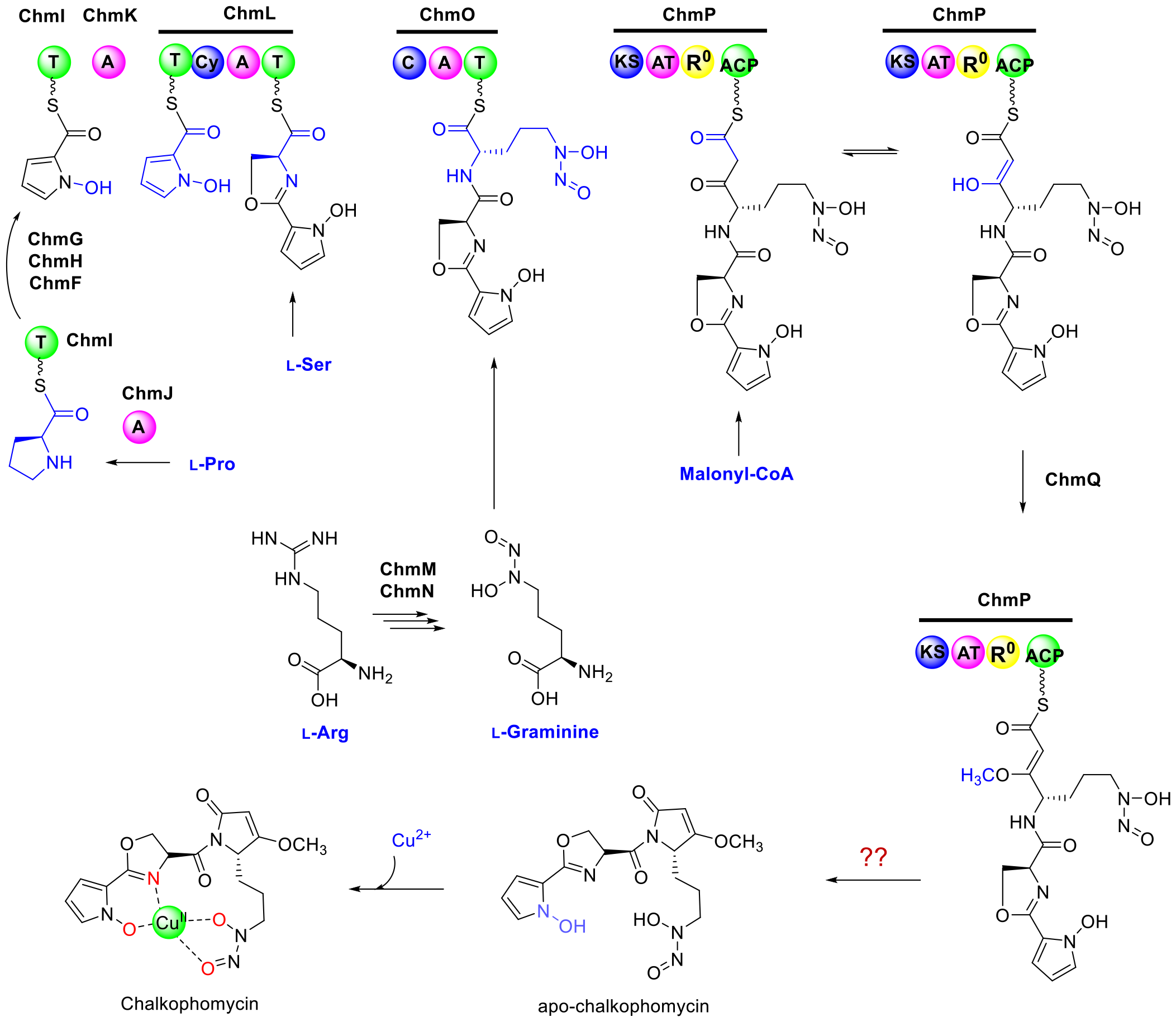
Chalkophomycin is proposed to be biosynthesized by a hybrid NRPS/PKS.

Biosynthesis of l-graminine monomer in chalkophomycin may involve the enzymatic action of ChmM and ChmN, since they shared 35% and 37% sequence identity with GrbE and GrbD, respectively. Synthetic l-graminine could restore the production of gramibactin in individual ΔGrbE and ΔGrbD mutants in *P. graminis* [11]. Therefore, l-graminine may be biosynthesized from l-Arg by ChmM and ChmN, followed by incorporation into chalkophomycin assembly line in S. sp. CB00271. It is also consistent with the recent observation by Bulter and co-workers that this unnatural amino acid l-graminine is derived from l-Arg in gramibactin biosynthesis by isotopic labeling studies [12].

#### 3.2.3. Chalkophomycin Biosynthesis by a Hybrid NRPS/PKS

After the conversion of l-Pro to pyrrolyl-*S*-ChmI catalyzed by ChmFGHIJ, a free-standing adenylation enzyme ChmK may mediate the transfer of the *N*-pyrrolyl intermediate to the first PCP domain on ChmL NRPS. ChmL is an NRPS with the characteristic PCP-Cy-A-PCP domain organization, in which the Cy (cyclization) domain is responsible for heterocycle formation in NRPS assembly lines, while the A domain in ChmL is predicted to activate l-cysteine. We proposed that this A domain is responsible for biosynthesis of the oxazoline moiety of chalkophomycin by loading l-serine to its cognate PCP, albeit further biochemical confirmation is needed. Both PCP domains of ChmO NRPSs and the discrete ChmI PCP with the signature motif of Gx(H/D)S, in which the Ser need be modified through the covalent attachment of the 4-phosphopantetheine group.

Following the ChmL NRPS is the ChmO protein with the characteristic NRPS condensation C-A-PCP domain organization, which may be responsible for activating and loading l-graminine to its cognate PCP domain, and the C domain may catalyze its condensation with the upstream dipeptidyl intermediate. Although the A domain in ChmO is predicted to activate l-*N*-hydroxyformyl ornithine, it is likely responsible for loading the unnatural l-graminine to chalkophomycin assembly line, since it also resembles the A domains for l-graminine activation in several NRPSs for the biosynthesis of megapolibactins, gladiobactin/plantaribactin with 30∼34% sequence identities (Figure S3) [11].

The *chmP* gene encodes a protein of 1414 amino acid residues containing one ketosynthase (KS), one malonyl-specific acyltransferase (AT), one domain with potential reduction function (named as R^0^) (amino acid residues 823–1299), and one acyl carrier protein (ACP). The KS is highly homologous to typical KSs from hybrid PKS/NRPS (Figure S3) [55–58], including EpoC for epothilone biosynthesis (Figure S4) [59]. The KSs in the hybrid NRPS/PKS would be responsible for transferring the dipeptidyl intermediate from the ChmO PCP domain, and catalyze the condensation with the incoming malonyl-ACP, mediated by its only AT domain. The AT-R^0^ didomain in ChmP shares 27% sequence identity with LnmG, a well-characterized, free-standing AT-oxidoreductase didomain protein for leinamycin biosynthesis [56]. AlphaFold prediction further reveals that ChmP_R^0^ domain might contain two Rossmann-like motifs (RLM) (Figure 4).

**Figure 4.**
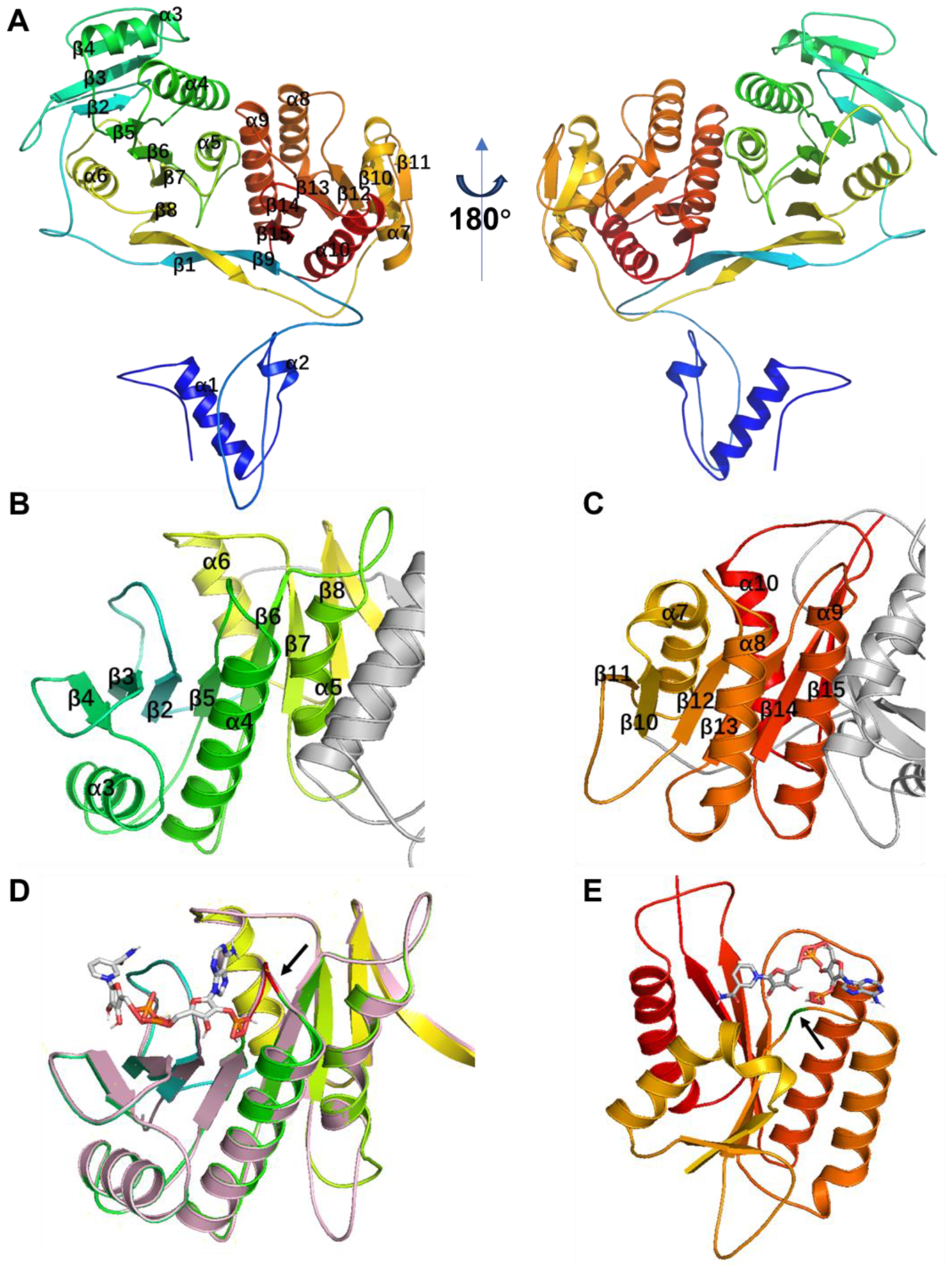
Structural analysis of ChmP_R^0^ domain with two putative Rossmann-like motifs (RLM).(A) AlphaFold2-generated model of the ChmP_R^0^ domain from *S*. sp. CB00271. (B–C) The RLM in the ChmP_R^0^ domain. (D–E) Docking analyses of the binding modes of NADP(H) to the ChmP_R^0^ domain from *S*. sp. CB00271 using AutoDock, respectively. The loop sequence marked in red is GSAAGPA in (D), while The loop sequence marked in green is GGG in (E). The RLM in (962–1180) is colored pink in (D).

We first employed Alphafold2 to predict the structure of the ChmP_R^0^ domain, which displays an alternating pattern of β-sheet-α-helix-β-sheet (Figure 4A). At the N- and C-terminus of ChmP_R^0^, there exists a double-wound three-layer α/β/α sandwich topology (Figure 4A–4C). The N-terminal region features a typical Rossmann fold with a central β sheet (strands β2–β7) arranged in the order 321456 (Figure 4B). Furthermore, this segment bears high similarity to the NADB region of the ChmP homologous protein from *Steptomyces* sp. MUN77 in both sequence and structure (Figures 4D and S5). The C-terminal includes a minimal RLM that crosses between the second and third strands in the order of 213456, and it is sandwiched between a layer of α helices (Figure 4C) [60]. These two RLMs are connected by a pair of antiparallel β-sheets.

The RLM usually possesses binding capability for diphosphate-containing cofactors such as NADP(H). The N-terminal turn of the first α-helix in RLM often binds to phosphate, and the gap between the first and third β-strands, β1 and β3, formed by the cross, could accommodate larger substrates or cofactors [61]. Accordingly, NADP was docked into both conservative pockets of RLM in ChmP_R^0^ domain using an AF2 prediction structure through AutoDock Vina [35]. The docking of NADP into the binding pocket of RML at the N-terminus of ChmP_R^0^ resulted in nine conformers with affinities ranging from –7.7 to –8.1 kcal/mol, while RML at the C-terminus of ChmP_R^0^ with affinities ranging from –6.8 to –7.1 kcal/mol. These docking analyses revealed that NADP can potentially bind to the cofactor binding pocket of both RLMs in the ChmP_R^0^ domain As shown in Figure 4D–4E, the structure with the highest binding free energy score was chosen for visualization. Additionally, each RLM in the ChmP_R^0^ domain contains a Gly-rich loop located in the C-terminal end of the β-sheet at termination of the crossover (indicated by the black arrow), which may become a part of RLM active site.

In the biosynthesis of cyclopiazonic acid in *Aspergillus* sp., a reductase-like R^*^ domain in the C-terminal of CpaS can carry out Dieckmann cyclization in a nonredox fashion [62]. Therefore, the R^0^ domain in ChmP may subtract one hydrogen from the ACP-tethered ketoamide, which leads to the formation of a resonance-stablized carbanion that undergoes O-methylation through ChmQ. Alternatively, the tethered ketoamide tautomerizes, and the resulting enol form could be methylated by ChmQ. Intriguingly, ChmP PKS lacks a type I thioesterase domain for acyl-ACP intermediate release upon completion of chain elongation in modular PKSs [63, 64]. The release of the peptidyl-acyl chain and the formation of an amide in the methoxypyrrolinone moiety may be executed non-enzymatically. During althiomycin biosynthesis in *M. xanthus* DK897 and an insect pathogen *Serratia marcescens* Db10, an iminopeptidase AlmF or a type II thioesterase Abl6 is proposed to play certain roles for chain release, respectively [13, 14]. Therefore, it remained to determine if the putative proline iminopeptidase ChmF may also facilitate chalkophomycin release.

#### 3.2.4. Regulatory and resistance for chalkophomycin biosynthesis

There are two putative regulatory genes in *chm* gene cluster, including *chmA* and *chmB*. ChmA gene encodes a regulatory LuxR family protein, which shows 35% sequence identity with RimR2, a recently identified positive pathway-situated regulator from *Streptomyces rimosus* M527 for rimocidin biosynthesis [65]. ChmB gene encodes a TetR/AcrR-like transcription regulator, which shows 32% sequence identity with SCO1718 from *Streptomyces coelicolor*. Four genes, *chmA’, chmC, chmD*, and *chmE*, could be identified within the *chm* gene cluster, which encode gene products to confer putative resistance to chalkophomycin. Both ChmA*’* and ChmC show 34% or 32% sequence identity of EfpA, a well-characterized multidrug efflux pump from *Mycobacterium tuberculosis* [66]. ChmD and ChmE encode a pair of ABC transporters, presumably to form an ATP binding cassette transporter complex responsible for chalkophomycin efflux.

### 3.3. Chalkophomycin-type gene clusters are wide-spread among phylogenetically diverse bacterial strains

Homologous *chm* gene clusters were first identified from GenBank, based on a cblaster search using genes including *chmA′* to *chmR*, and Blastn search using *chmP* as a query sequence; after manual removal of identical gene clusters, this search resulted in 39 homologous *chm* gene clusters (Figure 5 and Figure S6). With the availability of 11,357 assembled genomes in the Natural Products Discovery Center at University of Florida Scripps Research [39], ChmP-based BlastP search further resulted in 209 candidate gene clusters from 196 bacterial genomes. Close examination of the ChmP homologs in these candidate clusters suggested that the gene clusters with an identity cutoff of 50% for its ChmP homolog would result in 77 *chm*-type gene clusters. Next, *chm* and a total of these 116 *chm*-type gene clusters were analyzed by BIG-SCAPE, and their gene cluster similarity network was constructed and visualized using Cytoscape (Figure 5A). The majority of these gene clusters were closely clustered to *chm* gene clusters with 101 members. The other three gene cluster families contain 11, 2, and 2 members, respectively.

**Figure 5.**
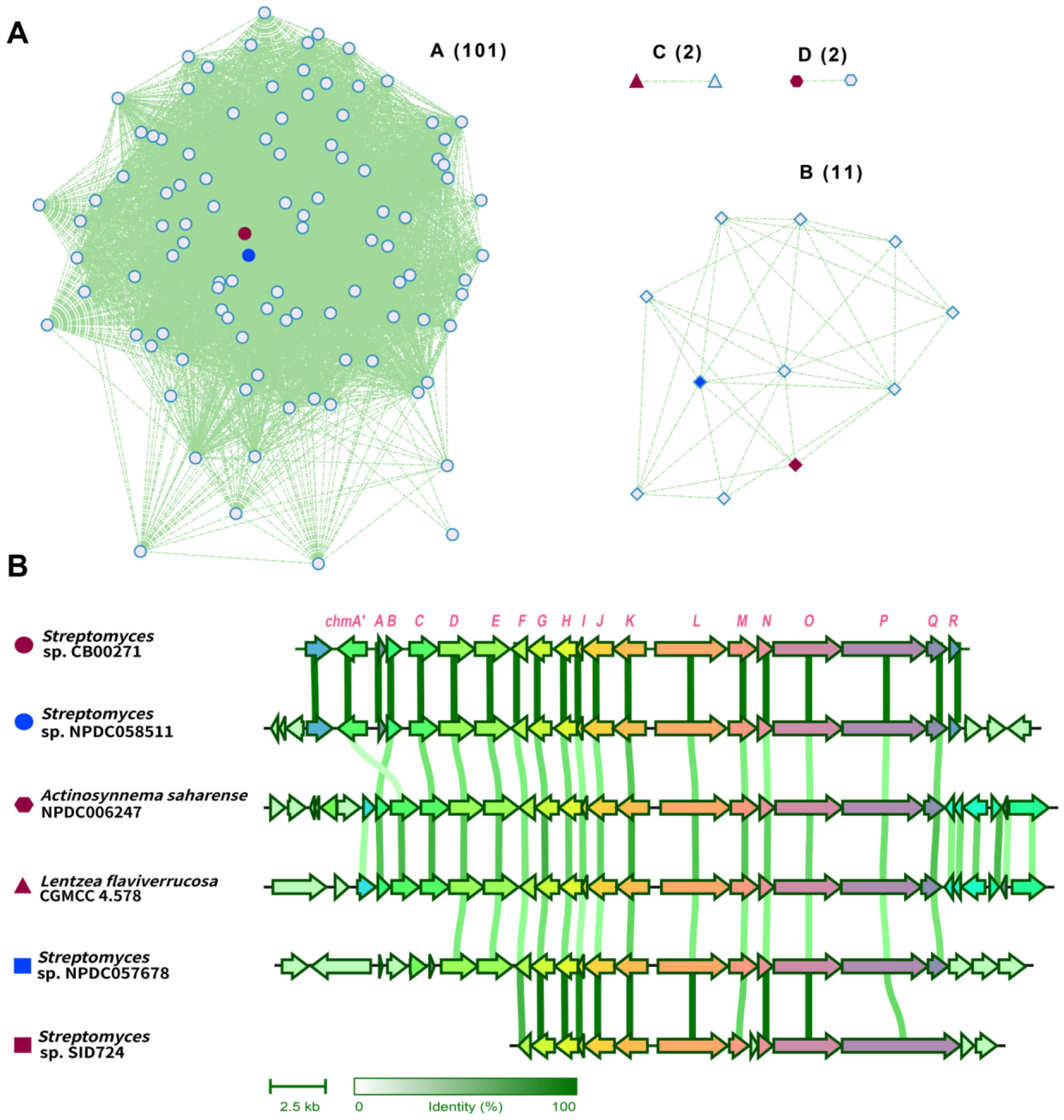
The global distribution of *chm* gene cluster in a wide range of actinomycetes.

Representative gene clusters from each cluster family were selected and aligned (Figure 5B). The *chm*-type gene cluster from *Streptomyces* sp. NPDC057678 is highly homologous to *chm* gene cluster from *S*. sp. CB00271, showing >95% sequence identity across *chmA’* to *chmR*. In both *chm*-type gene clusters in rare actinomycetes, e.g., *Actinosynnema saharense* NPDC006247 and *Lentzea flaviverrucosa* CGMCC4.578, the multi-drug transporter EmrB/QacA is instead positioned downstream of two genes encoding a TetR transcription regulator and a molybdenum co-factor sulfurase C-terminal domain protein, respectively. Furthermore, there are several notable differences of the *chm*-type gene cluster from *Streptomyces* sp. SID724, including (a) a significantly larger PKS (1914 a.a) with a KS-AT-dehydratase-reductase domain organization, (b) an additional small cupin domain-containing protein, and (c) the lack of an *O*-methyltransferase. Therefore, a new chalkophore with distinct structures may be produced in this specific strain. Taken together, these analyses suggest that *chm*-type gene clusters are widely distributed in actinomycetes, especially in *Streptomyces*. It remained to determine if the presence of these genes clusters conveys certain survival advantage for the host strain, since the essential cupric ions may be assimilated to the host through the produced chalkophores. Alternatively, chalkophomycin was also toxic to some tested bacteria, and the producing strain may also have advantage over its competitor in the surrounding environment.

## 4. Discussion

In this study, the biosynthetic gene cluster for chalkopomycin was identified from *S*. sp. CB00271, revealing an unusual hybrid NRPS/PKS with an atypical R^0^ domain in a PKS module, which might contain two RLMs for the binding of two NADP(H) co-factors. In addition, over 100 homologous *chm* gene clusters were discovered from public databases, suggesting the widespread of this gene cluster.

Chalkophores are relatively rare in nature comparing to siderophores, and the prototypical chalkophores methanobactins are biosynthesized ribosomally and then undergo extensively morphing [17–19]. In contrast, chalkophomycin biosynthesis follows the assembly model of hybrid NRPS/PKS. In most characterized NRPS/PKSs, these mega-synthetases/synthases often encompass multiple modules, with typical domain organization of C-A-PCP in a NRPS module and KS-AT-(KR-DH)-ACP in a PKS module. There are extensive intra- or intermodular communications mediated by conserved linkers responsible for efficient transfer of peptidyl- or acyl-intermediates. The assembly line models were studied using several model systems, e.g. DEBS for PKSs, and surfactin and tyrocidine for NRPSs [67, 68]. However, the complex module organization of these huge mega-synthases poses formidable challenges to isolate, purify, and characterize them biochemically and structurally [69]. ChmL/ChmO/ChmP are all single module protein, and thus provide a rare opportunity to study their inter-modular communications. In particular, how the peptidyl-PCP from ChmL was transferred to ChmP KS. In addition, the availability of these *chm*-type gene clusters may provide additional possibilities to study chalkophomycin biosynthesis in an evolutionary context, as we have observed in the genome neighborhood analyses (Figure 5). These efforts may lead to the discovery of new enzymology for chalkophore biosynthesis and potential for a synthetic biology approach to discover and engineer chalkophores.

Although copper homeostasis is essential for most life on earth, most actinomycetes are soil-dwelling bacteria. The identification of over 100 homologous *chm*-type gene clusters in not only *Streptomyces* species, but also in rare actinomycetes, i.e. *Actinosynnema saharense* and *Lentzea flaviverrucosa*, not only suggest the horizonal gene transfer of the gene clusters, but also imply the important role of chalkophomycin and alike for these micro-organisms. Considering the discovery of several chalkophores from fungi and bacterial pathogens [17], understanding their physiological roles may be instrumental to study the role of copper ions in living organisms, including Homo sapiens [29].

## Supplementary Materials

Table S1: Plasmids and strains used in this study; Table S2: Primers in this study; Figure S1: Identification of three sets of homologous proteins of GrbED from *S*. sp. CB00271, using query sequences GrbE (WP_006051176.1) (A) and GrbD (WP_006051175.1) (B); Figure S2: Phylogenetic analysis of ChmQ with other known methyltransferases; Figure S3: Alignment of the adenylation domain of ChmL predicted for activation of L-graminine with other L-gramininespecific adenylation domains from megapolibactins and gladiobactin/plantaribactin; Figure S4: Phylogenetic analysis of the KS domain of ChmP with other known KSs using NaPDoS2 webtool; Figure S5: Sequence alignment of the R^0^ domain of ChmL from *S*. sp. CB00271 and *Streptomyces* sp. MNU77; Figure S6: Phylogenetic analysis of chm gene cluster from *S*. sp. CB00271 and 116 identified chm-type gene clusters from the public databases.

## Author Contributions

Conceptualization, Y.H.; methodology, L.Y., L.W.Y., L.C., B.G., M.L.; software, L.C. and Y.L.; validation, L.Y. and L.W.Y.; formal analysis, L.Y., L.W.Y., L.C.; investigation, L.Y., L.W.Y., L.C.; resources, L.Y., Y.D., and Y.H.; data curation, L.Y., L.W.Y., L.C.; writing—original draft preparation, L.Y., L.Y., L.C. and Y.H.; writing—review and editing, L.Y., L.Y., L.C. and Y.H.; visualization, Y.L. and L.C.; supervision, X.Z. and Y.H.; project administration, Y.H.; funding acquisition, L.Y., Y.D., and Y.H.

## Data Availability Statement

Data are contained within the article and Supplementary Materials.

