## Supporting Information for "Chalkophomycin Biosynthesis Revealing Unique Enzyme Architecture for a Hybrid Nonribosomal Peptide Synthetase and Polyketide Synthase"

[#]Equal contribution.

### Index

**Table S1**. Plasmids and strains used in this study. ........ S3
**Table S2**. Primers in this study. ........ S4
**Figure S1**. Identification of three sets of homologous proteins of GrbED from S. sp. CB00271, using query sequences GrbE (WP_006051176.1) (A) and GrbD (WP_006051175.1) (B). ........ S5
**Figure S2**. Phylogenetic analysis of ChmQ with other known methyltransferases. ........ S6
**Figure S3**. Alignment of the adenylation domain of ChmL predicted for activation of L-graminine with other L-graminine-specific adenylation domains from megapolibactins and gladiobactin/plantaribactin. ........ S7
**Figure S4**. Phylogenetic analysis of the KS domain of ChmP with other known KSs using NaPDoS2 webtool. ........ S8
**Figure S5**. Sequence alignment of ChmL proteins from *S*. sp. CB00271 and *Streptomyces* sp. MNU77 with $R^0$ domain highlighted in red. ........ S9
**Figure S6**. Phylogenetic analysis of *chm* gene cluster from *S*. sp. CB00271 and 116 identified *chm*-type gene clusters from the public databases. ........ S10
**References**. . ........ S11

**Table S1**. Plasmids and strains used in this study.

| Strains/plasmids | Descriptions | Reference/ Source |
|---|---|---|
| *E. coli* strains | | |
| DH5α | General cloning | Commercial source |
| S17-1 | Intergeneric conjugal transfer | Kieser T, 2000 |
| *Streptomyces* strains | | |
| CB00271 | Producing strain of chalkophomycin | This study |
| CB00271::Δ*chmO* | *ChmO* gene replacement mutant of CB00271 | This study |
| Plasmids | | |
| pOJ260 | *E. coli* vector, nonreplicating in *Streptomyces*, $Am^R$ | Kieser T, 2000 |
| pXY5001 | pOJ260-based plasmid for *chmO* inactivation, $Am^R$ | This study |

**Table S2**. Primers in this study.

| Primers | Sequences (5'-3') | Function |
|---|---|---|
| NRPS1.1-Up-F* | GCGGCCGCGGATCCTCTAGAGC GGCACAGGGCGAG | Construction of knockout plasmid for *chmO* |
| NRPS1.1-Up-R* | GAATGTGAACACACCTCGCGCA CCCC | |
| NRPS1.1-Tsr-F* | TGCGCGAGGTGTGTTCACATTC GAACGGTCTCTGC | |
| NRPS1.1-Tsr-R* | GGGCCAACTGATTTATCGGTTG GCCGCGAGA | |
| NRPS1.1-Dn-F* | GCCAACCGATAAATCAGTTGGC CCGCCG | |
| NRPS1.1-Dn-R* | ACGACGGCCAGTGCCAAGCTT CCTGCCGACGAACTCGA | |
| pYZNRPS1.1F | CCGCGCTTCGTCGATCTG | confirmation of mutant Δ*chmO* |
| pYZNRPS1.1R | TCCAGCAGCACCCCGAC | |

* The red-labeled nucleic acid sequence is the overlapping region with pOJ260 and the homology arm.

**Figure S1**. Identification of three sets of homologous proteins of GrbED from *S*. sp. CB00271, using query sequences GrbE (WP_006051176.1) (A) and GrbD (WP_006051175.1) (B).

**A**

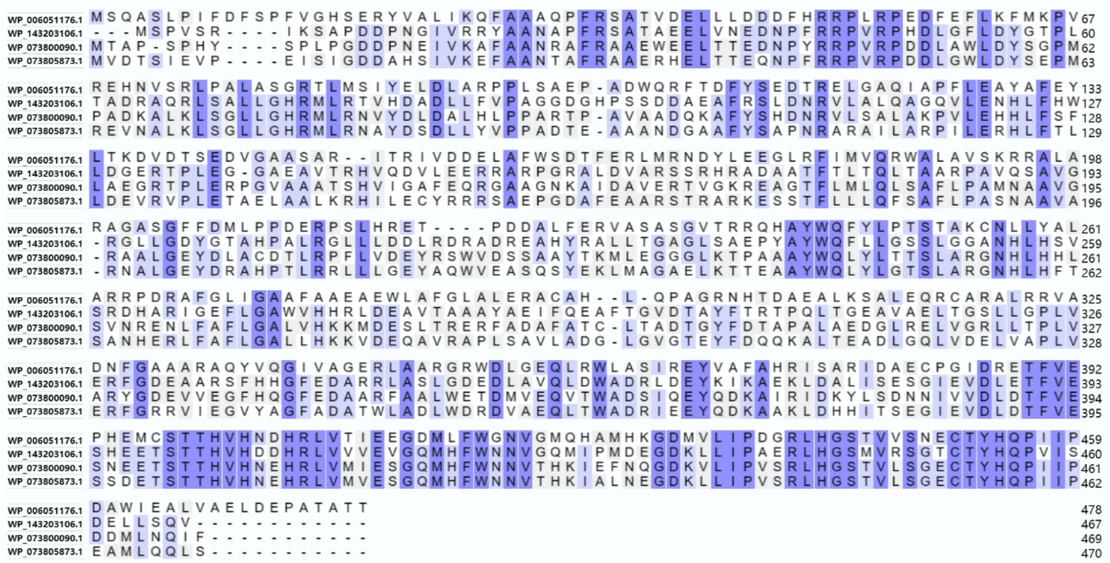

**B**

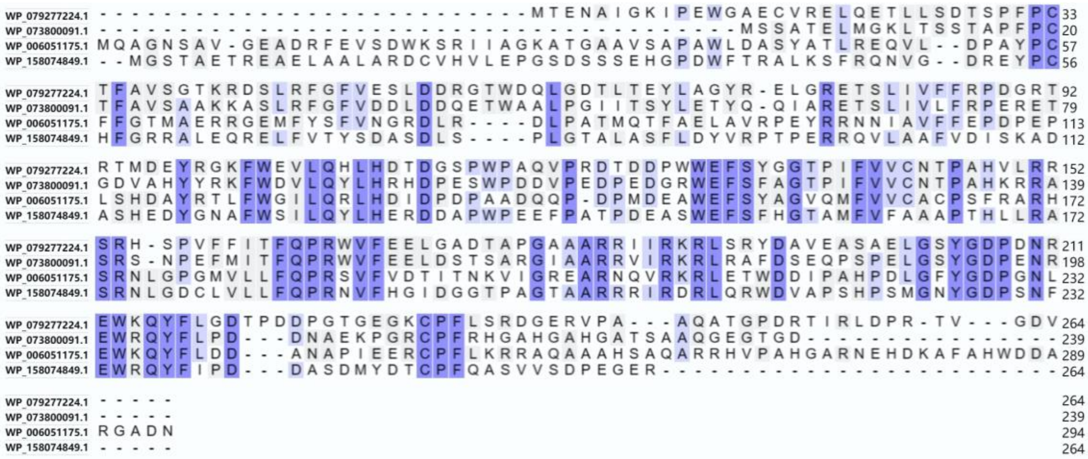

**Figure S2**. Phylogenetic analysis of ChmQ with other known methyltransferases.

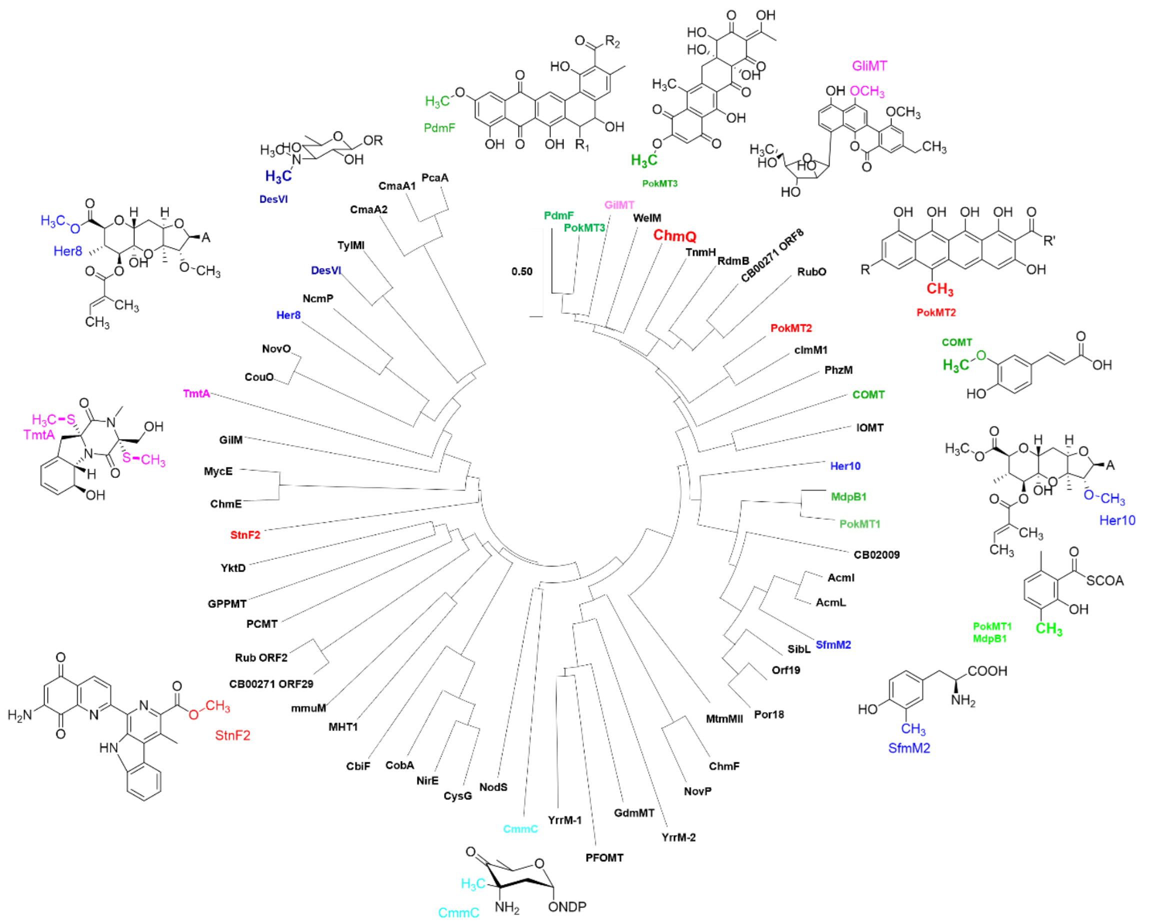

**Figure S3**. Alignment of the adenylation domain of ChmL predicted for activation of L-graminine with other L-graminine-specific adenylation domains from megapolibactins and gladiobactin/plantaribactin.

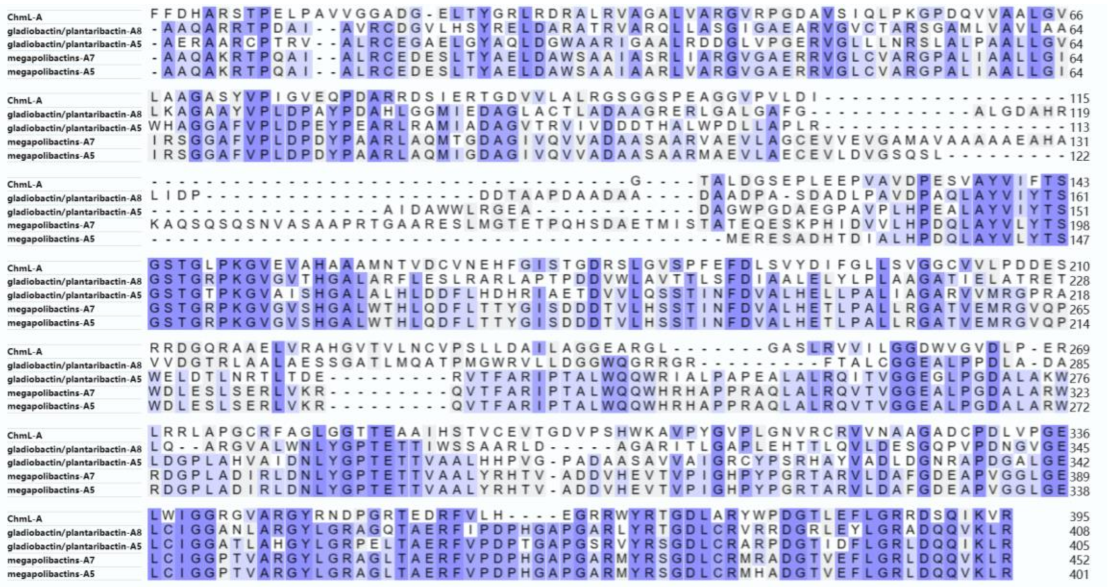

**Figure S4**. Phylogenetic analysis of the KS domain of ChmP with other known KSs using NaPDoS2 webtool.

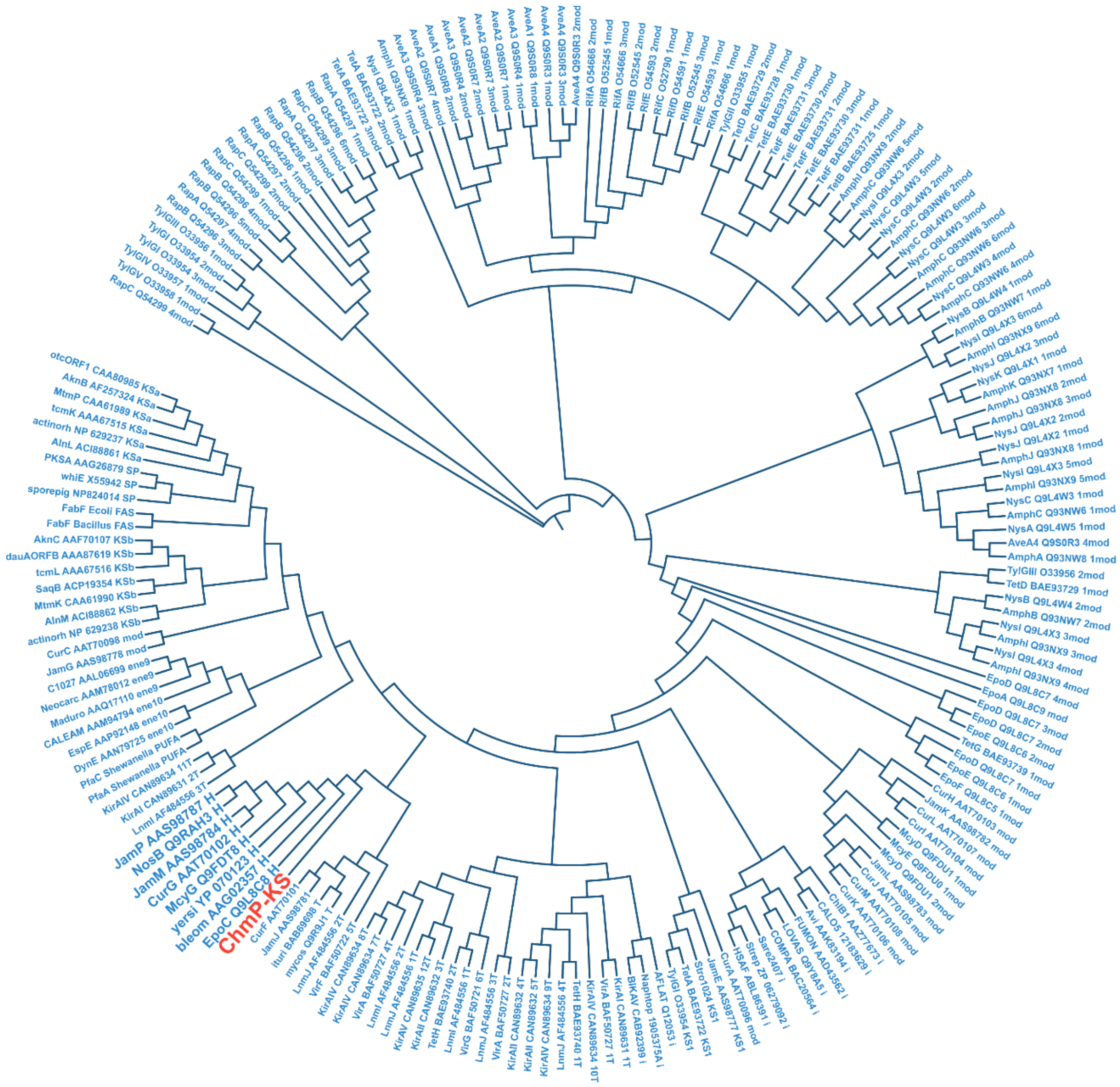

**Figure S5**. Sequence alignment of ChmL proteins from *S.* sp. CB00271 and *Streptomyces* sp. MNU77 with R$^0$ domain highlighted in red.

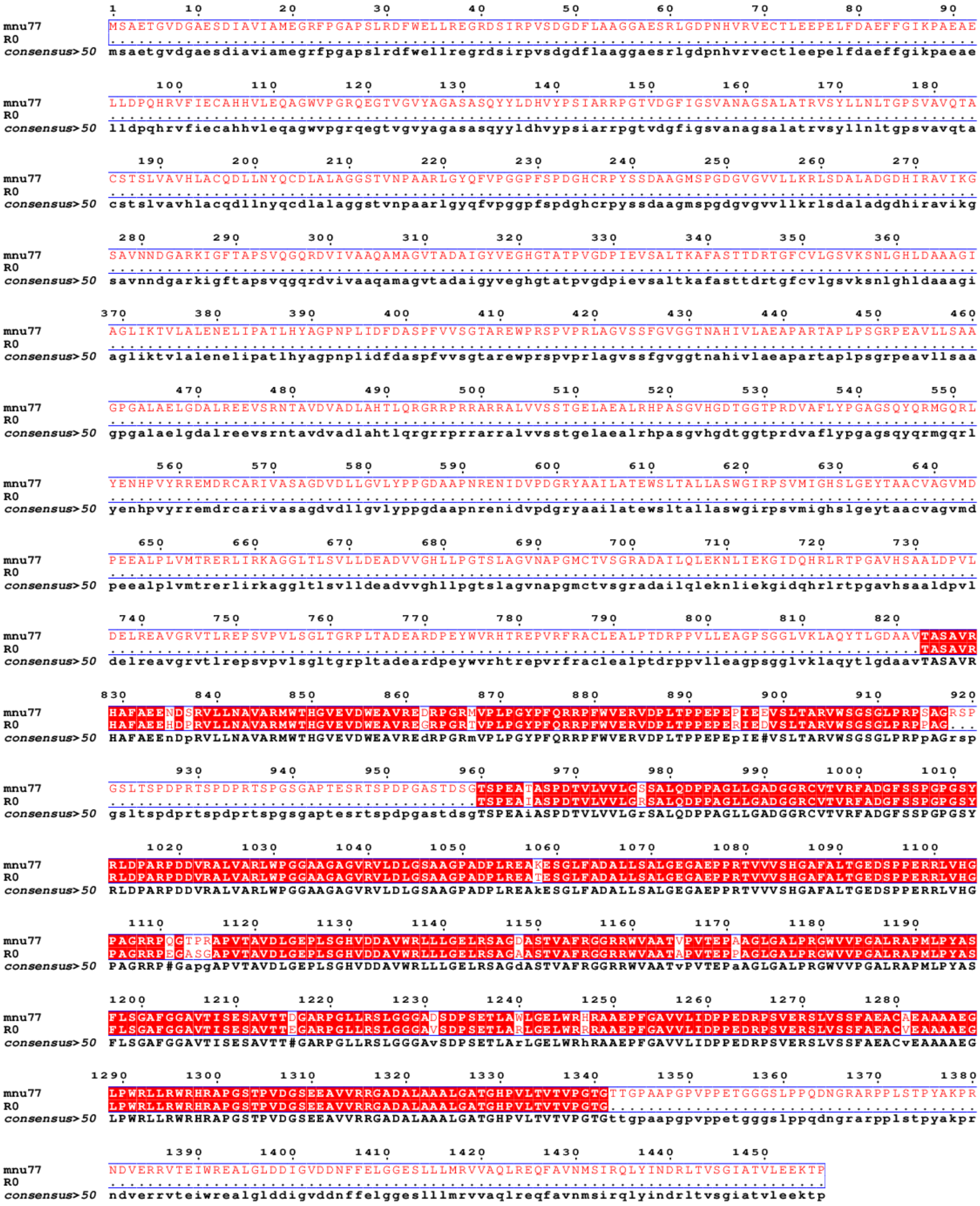

**Figure S6**. Phylogenetic analysis of *chm* gene cluster from *S*. sp. CB00271 and 116 identified *chm*-type gene clusters from the public databases.

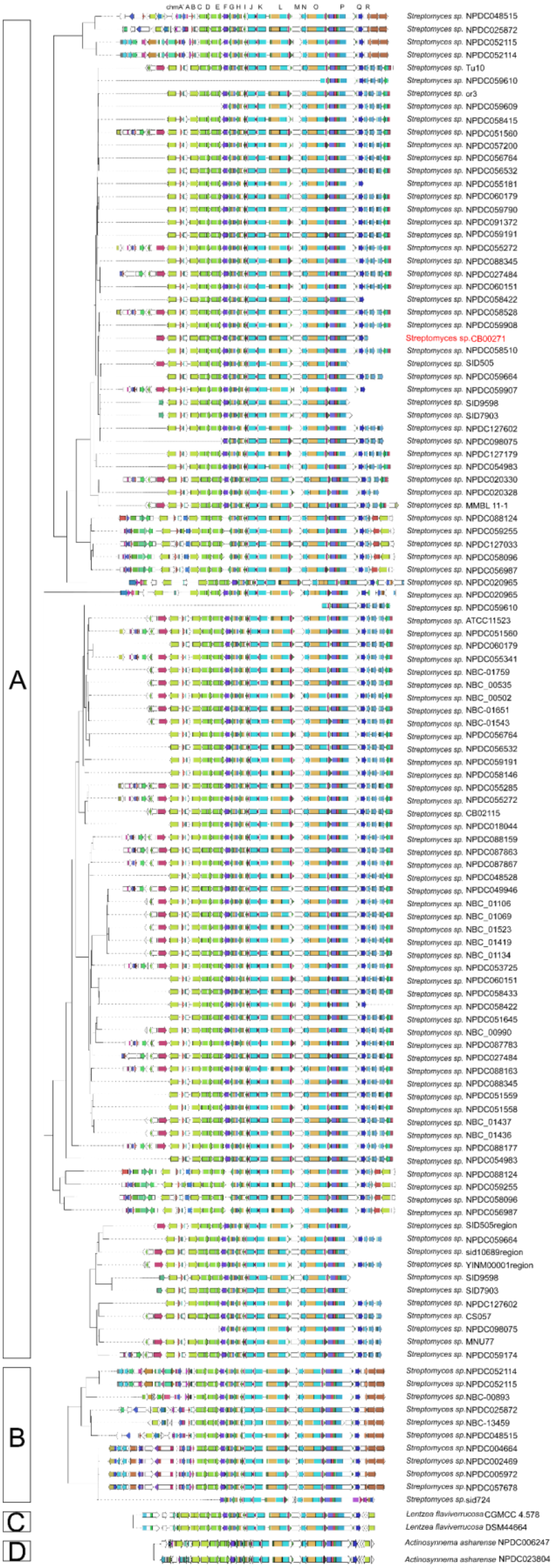

**References**.

1. Kieser, T.; Bibb. M.J.; Buttner, M.J.; Chater, K.F.; Hopwood D.A. (2000). Practical *Streptomyces* genetics. *John Innes Foundation, Norwich, United Kingdom.*
